# Deciphering fitness trade-offs in metabolite exchange at the origin of a bacterial cross-feeding community

**DOI:** 10.1101/2023.11.13.566325

**Authors:** Dongxuan Zhu, Samraat Pawar

## Abstract

Obligate cross-feeding is a common type of interaction among microbial communities, yet gaps persist in understanding its maintenance and limit practical applications. In particular, little is known about how contextualised metabolite exchange intensities affect community fitness, despite their influential role in shaping interdependencies^1^, diversities ^2^, and lifespan ^3^ of cross-feeding community members.

Here I computationally test how amino acids isoleucine (*ile*) and lysine (*lys*) exchange intensities affect individual and community fitness of a two-strain auxotroph cross-feeding community. I innovatively integrated metabolite exchange intensities and multi-strain growth using Flux Balance Analysis (FBA) and evolutionary game theory, and showed that crossfeeding communities have the highest fitness when the metabolite exchange intensity is slightly above individual amino acid uptake demands, stimulated by cheaters’ presence. Using FBA with different metabolite uptake / secretion combinations, I discovered the individual amino acid demands are linearly correlated with sole carbon source (glucose) availability. Additionally, as cheating mutants emerge, costly exchange intensities can be better sustained when accumulated shared metabolites are accessible.

This is the first study linking metabolite exchange intensities and cross-feeding community fitness considering all known metabolic reactions of a bacterial strain. The fittest metabolite exchange range and its relationships with glucose and shared metabolite availability shed lights on resilient microbial community engineering as well as metabolite exchange parameter constraints for multi-species population / metabolism models ^4–6^.

## 1 Introduction

Obligate metabolite secretion and exchange are widely observed in natural microbial communities^7,8^. In particular, obligate cross-feeding - the process where microbial species exchange essential nutrients - plays a key role in improving microbial community resilience and metabolic efficiency^5,9–11^. Therefore, engineered cross-feeding communities have wide application potentials from plastic waste degradation^12^ to pharmaceutical production^13^. However, current engineered microbial communities cannot sustain their performance due to adverse events such as mutations and environmental changes^14,15^. Particularly, it is not yet clear how to maintain fitness of the cross-feeding community members when cheating mutants emerge.

To date, accumulated evidence suggests that metabolite exchange intensity plays an important role in shaping cross-feeding community dynamics^2–4^. A theoretical study by ^4^ demonstrates that increasing metabolite exchange intensity significantly enhances coalescence success between two microbial communities. Furthermore, several empirical and theoretical studies find that an increase in metabolite exchange tend to trigger the co-evolution process where interdependency emerges^1,16,17^. Therefore, a further understanding on how metabolite exchange intensities affect cross-feeding community fitness is an intriguing goal to achieve.

Unfortunately, most relevant studies to date do not integrate evolutionary dynamics with ecophysiological consequences of different metabolite exchange rates. On one hand, both empirical and theoretical evolutionary studies have explored co-evolutionary dynamics originating from imposed obligate cross-feeding^1,16,18^, but they lack a metabolic linkage to fitness costs and benefits stemming from metabolite exchange. On the other hand, theoretical studies that do consider realistic metabolic reactions make arbitrary assumptions in evolution, ignoring the potential of cheating mutants or assuming fast fixation between mutations^19,20^.

For a more comprehensive study integrating both valid evolutionary context and ecophysiological properties, evolutionary game theory in conjunction with the Flux Balance Analysis (FBA) method could be a promising approach. Different from typical evolutionary dynamics where fitness of one phenotype depends on its own reproduction success under certain environmental conditions, evolutionary game theory considers interactions among all phenotypes along with frequencies of these phenotypes^21^. This is especially helpful when studying cross-feeding interactions, where microbes tend to physically attach to one another (e.g. via nanotubes) and therefore rely on partner frequencies for reciprocal benefits^22^. Another key concept in evolutionary game theory is the notion of evolutionarily stable state (ESS), where a community cannot be invaded by any mutants^21^. Survival in such invasion experiments can therefore serve as a benchmark for determining the optimal choice of metabolite exchange strategies^23^. With eco-evolutionary framework set up by evolutionary game theory, the next step is to ground individual fitness with realistic metabolic information. FBA is a powerful tool since it takes into account all known metabolic reactions of an organism^24^. Its prediction has been proven to have good agreement with empirical data for both single species^25^ and two-to three-species communities^26^. Given its accurate depiction of growth based on metabolic functional traits, FBA results can effectively represent the net benefits of different encounters in evolutionary game theory.

The most comprehensive combination of FBA with evolutionary game theory to date was introduced by. ^27^ In detail, they constructed two auxotroph *E*.*coli* strains that fed on each other and forced to secrete certain amount of amino acid, mediated by both empirical data of maximum amino acid production rates^28^ and number of consumers in each cell-cell encounter. They then varied the chosen amino acids and secretion levels in a series of invasion experiments involving resident wildtype and auxotroph strains, as well as secretion-free invaders. Similar to findings from, ^1^ a cross-feeding community seems to be close to an evolutionarily stable state when the metabolite secretion intensity is approximately 10-30% of the maximum level. However, these experiments assume constant population size with constant benefits in growth simulation, ignoring the stochastic growth process and the effect of metabolite accumulation. The mutant strain with deficiency in synthesising all required amino acids is also too dependent on the existence of the wildtype strain. As these factors might affect invasion outcome^29^, improvements need to be made.

To build upon existing research and address these gaps, I aim to answer the following questions with this study:

1. What are the fitness trade-offs for secreting and uptaking growth-limiting metabolites, and how they are affected by the media conditions, especially the depletion of glucose and accumulation of shared metabolites?
2. With various metabolite exchange intensities, under what intensities will cross-feeding become the fittest strategy to maintain, and why?

## 2 Methods

### Rationale

The primary aim of my study is to understand how metabolite exchange intensities affect the fitness of engineered cross-feeding communities, under the threat of cheating mutants. In order to estimate community fitness, I adopted the invasion tests and the notion of evolutionarily stable states from evolutionary game theory. Two cross-feeding auxotroph strains as the ancestors and two cheating mutant strains as the invaders are constructed to imitate different expression strategies in metabolite synthesis and secretion^30^. In the invasion tests, I forced each cross-feeding auxotroph to secrete the same rate of shared metabolites in one invasion event to imitate one metabolite exchange strategy, and vary metabolite exchange strategies by invasion events (Fig. 1). A resident cross-feeding community persist such an invasion will be close to evolutionarily stable state as it outperforms the fittest mutant.

**Figure 1:**
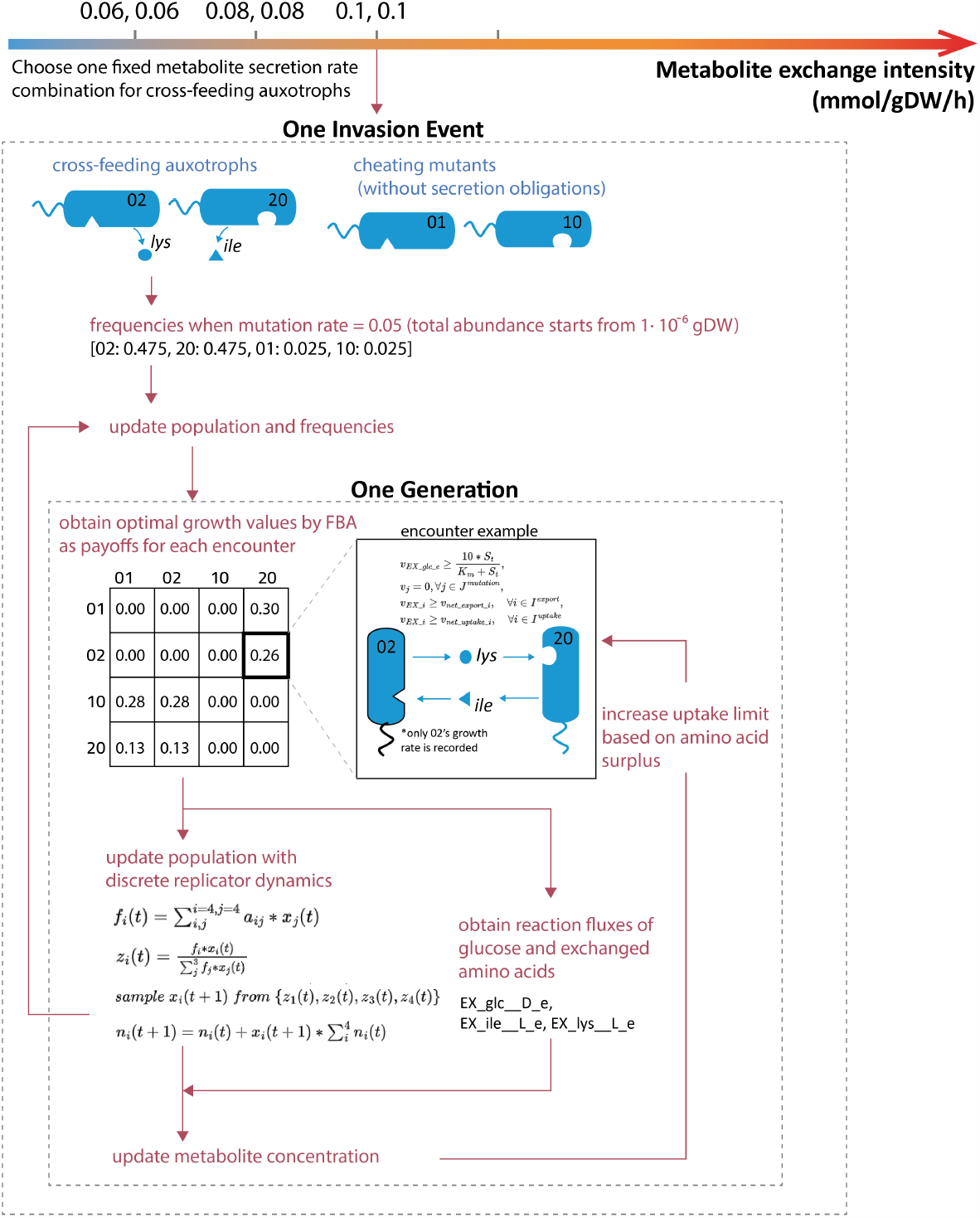
Flow chart showing the modelling framework of invasion experiments. For each generation updates, growth rates are calculated by FBA and recorded in payoff matrix for the left side strains. Formula in encounter example are part of the FBA formula related to key metabolites.

The growth simulation model in invasion tests follows the principles of discrete replicator dynamics in evolutionary game theory, except for its combination with FBA and consideration of actual population abundance (Fig. 1). The former is introduced so that extra nutrient concentration can be involved in fitness updates, and the latter is necessary for computing metabolite concentration.

In addition to invasion tests, I also simulate a series of one-generational monoculture growth of the two cross-feeding strains at different amino acid uptake and secretion rates. This step helps me to understand strains’ differences in intrinsic growth rates.

In terms of the growth-limiting metabolites they exchange, I chose amino acids as focus for their essential roles for growth and prevalence in natural cross-feeding communities^31^. I chose a minimal medium with limited glucose supply to avoid overflow metabolism^32^.

### Media and culture conditions

The media component design aims to support minimal growth for auxotroph strains. Adapted from the Long-Term Evolution Experiment (LTEE) of E. coli^33^ and its in-silico reproduction^34^, I conducted simulations in this study using a 1L batch reactor with a 10 mmol/L supply of DGlucose. I then defined other media compounds based on the M9 minimal media database from CarveMe ^35^. I configure in-silico uptake rates of glucose and oxygen as 10 mmol/gDW/h and 2 mmol/gDW/h, respectively. In addition, I set the ATP maintenance rate at exactly 8.39 mmol/gDW/h based on existing observations^36^. I set uptake rates for all other compounds to 1000 mmol/gDW/h in order to elevate corresponding growth constraints. I also set the maximum secretion rate (upper bound fluxes) for all reactions to 1000 mmol/gDW/h.

To accurately predict strain behaviors in-silico, I downloaded a manually curated genome-scale metabolic model of E. coli iJO1366 from BiGG Models (http://bigg.ucsd.edu). Subsequently, I imported the downloaded SBML file as a COBRA model using COBRApy. Then, I configured the initial media for growth simulation by setting the lower bound for exchange reactions of the COBRA model.

### Construction of auxotroph strains with different metabolite exchange intensities

To ensure the inter-dependency of the cross-feeding community in invasion experiments, I constructed two auxotroph strains by knocking out their genes that regulate complementary amino acid synthesis processes. To ensure auxotroph strains in a small community can cross-feed each other without large difference between growth rates before invasion, I chose the amino acids L-isoleucine (*ile*) and L-lysine (*lys*) as my focus due to their similar growth costs and benefits.

I then knocked out genes regulating the synthesis processes for *ile* and *lys* with COBRApy, and obtain two auxotroph genotypes, 02 and 20. Strain 02 is an auxotroph for *ile* (annotated 0), but synthesises and secretes *lys* (annotated 2). Conversely, strain 20 is an auxotroph for *lys*, but synthesises and secretes *ile*. I aslo introduce two extra phenotypes, 01 and 10, based on the same genome information as 02 and 20, but I lifted their secretion rate constraints for *lys* and *ile*. Except for these lifted constraints, the net secretion or uptake rate of all strains is constrained based on the following formula:

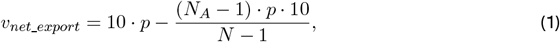

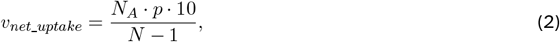

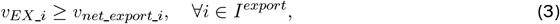

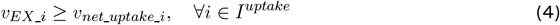

Where *N* denotes total number of genotypes in a given encounter, *N*_*A*_ denotes number of genotypes that secret amino acid A in this encounter, and 10 is the constant representing maximum export flux of all types of amino acid based on observation by. ^27^ Parameter *p* represents the proportion of amino acid secreted to extracellular environment (i.e. become public goods), which is a key variable representing strain’s secretion capability for specific amino acid. In addition, *v*_*EX*_*i*_ represents flux of exchange reactions for amino acid i, and *I*^*uptake*^ and *I*^*export*^ respectively denote available amino acid for uptake or export for each strain. Because physical attachments such as nanotubes are common when bacteria cross-feed amino acids^22,37^, metabolite loss via diffusion is not considered in the simulations. Maximum uptake rate of a recipient strain is almost equal to minimum secretion rate of a donor strain - with only addition of metabolites from shared medium.

### One-generational growth simulation with different metabolite exchange intensities

To have preliminary understanding of the costs and benefits of metabolite exchange, I conducted a series of one-generational growth simulation with two auxotroph strains at different metabolite secretion and uptake levels. Each combination is simulated with 21 glucose supply levels ranging from 5 to 15 mmol/gDW/h to include the impact of glucose depletion into consideration.

Each simulation is fulfilled by FBA using COBRApy under python 3.8 environment. Based on the media condition and auxotroph genotypes described above, I constrained the minimum secretion rate of their possessed amino acids, and fixed the uptake rate of their essential amino acids. Growth rates are then calculated by optimizing their biomass yield flux.

Monoculture experiments are simulated based on biomass yield responses to metabolite supplies. These experiments assume auxotrophs consume glucose and secrete amino acids at the same rates as in invasion experiments at each time step. Auxotrophs are constantly supplied with the same amount of complimentary amino acids as the given metabolite exchange intensities.

### Invasion experiments with different metabolite exchange intensities

In evolutionary game theory, a community is considered as evolutionary stable if it can resist invasions from any mutants. To mimic this situation and test whether a cross-feeding community is an evolutionary stable community, I constructed a cross-feeding community [02, 20] with its secretion-free invaders [01, 10] using the strains described in section 2.2. Invasion experiments are then conducted with cross-feeders at different metabolite secretion levels, variating within feasible range.

Each experiment starts from the frequency at [0.475, 0.475, 0.025, 0.025] for strain [02, 20, 01, 10]. Abundance of 02 and 20 starts from 4.5×10^*−*7^ gDW so that glucose consumption starts from low rate. For each experiment, generation update is completed in following steps:

1. Examine of all interactions between phenotypes and evaluate respective benefits using growth yield values obtained by FBA. These benefit values are stored associated with participants’ phenotypes as payoff matrix. It is then utilised for fitness evaluation.
2. Update metabolite concentration in each growing time step.
3. Update population in each growing time step. I use 20min as the interval for growth as it is the typical doubling time for *E*.*coli*. I use discrete replicator dynamics for growth simulation, which is adapted from an evolutionary modelling study by. ^38^ It needs to be noted here that only growth and fixation phases are simulated here, excluding death phase.

### Compute dynamic payoff values for each strain

For each strain in step 1’s interactions, FBA problem is solved based on metabolite constraints. The objective for FBA is all set to optimize individual biomass growth with the problem formulated as follows:

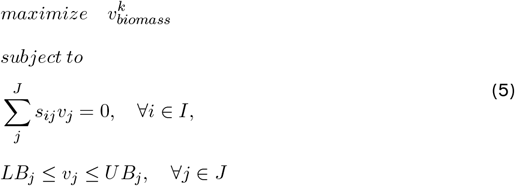

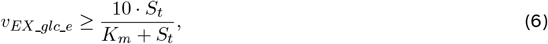

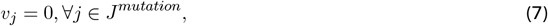

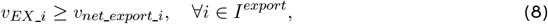

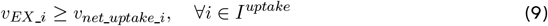

Where *s*_*ij*_ refers to stoichiometry matrix for the strain, *J* represents the set of all reactions, and *J*^*mutation*^ refers to reactions to be knocked out for mutant strains. Uptake of glucose from medium follows Michaelis-Menten function to mimic concentration-dependent uptake, where 10 is the maximum uptake rate of glucose and *S*_*t*_ is the concentration at time step *t. K*_*m*_ referes to the half-saturation constant of glucose transporter in mmol/l, which takes value at 0.02 according to empirical studies^39^.

### Update concentration of shared metabolites

To maximize biomass growth under constraints, strain might secrete more metabolites than required^40^. At the same time, these redundant public goods have an significant impact on crossfeeding interactions.^29^ Therefore, I record these surpluses and add them to relevant amino acid uptake constraints (Eq 9). Firstly, strain frequency is updated based on previous payoff matrix; Secondly, exchange reaction fluxes of chosen amino acids for each strain are stored and summed, resulting in net flux surpluses; Lastly, these surpluses are added to corresponding uptake reactions of each strain in new FBA calculation (1).

### Update population with Discrete Replicator Dynamics

In order to allow prompt payoff value update and integrate FBA in the process, I adapt discrete replicator dynamic model from^38^ to model population growth of different strains.

Let *x*_1_, *x*_2_,*x*_3_ and *x*_4_ symbolize the relative proportions of four strains, and *f*_*i*_(*x*_1_, *x*_2_, *x*_3_, *x*_4_) for *i* = 1,2,3,4 represent functions that define their frequency-dependent fitness. The ecological dynamics within finite populations and discrete time is defined by:

i. Calculate fitness function: 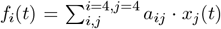, where *a*_*ij*_ refers to benefit gained by strain i when encountering strain j.
ii. Summarize average fitness value for strain i: 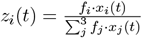
iii. sample new frequency *x*_*i*_(*t*+1) from the multinomial distribution denoted by *{z*_1_(*t*), *z*_2_(*t*), *z*_3_(*t*), *z*_4_(*t*)*}* with *N* individuals in total. *N* use an arbitrary number 1.00×10^6^ to ensure sampling resolution;
iv. Update population abundance: 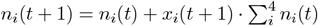

This serves as a basic model of ecological dynamics involving pairwise interactions, relying on the averaging of outcomes from all feasible sets of interacting strains.

## 3 Results

### Exchanged amino acids have growth cost and benefit thresholds mediated by glucose availability

Bacterial auxotrophs can adapt amino acid uptake and secretion rates to various metabolic environment ^31,41^. To investigate what metabolite exchange rates induce the highest individual fitness of cross-feeding auxotrophs, I simulated their instantaneous biomass yields over one generation with different combinations of amino acid uptake and secretion rates. Simulation results show significant piecewise growth costs and benefits by increasing amino acid uptakes or secretions, suggesting the existence of metabolite exchange thresholds that mediate bacterial auxotroph fitness with more than one mechanisms (Fig. 2).

**Figure 2:**
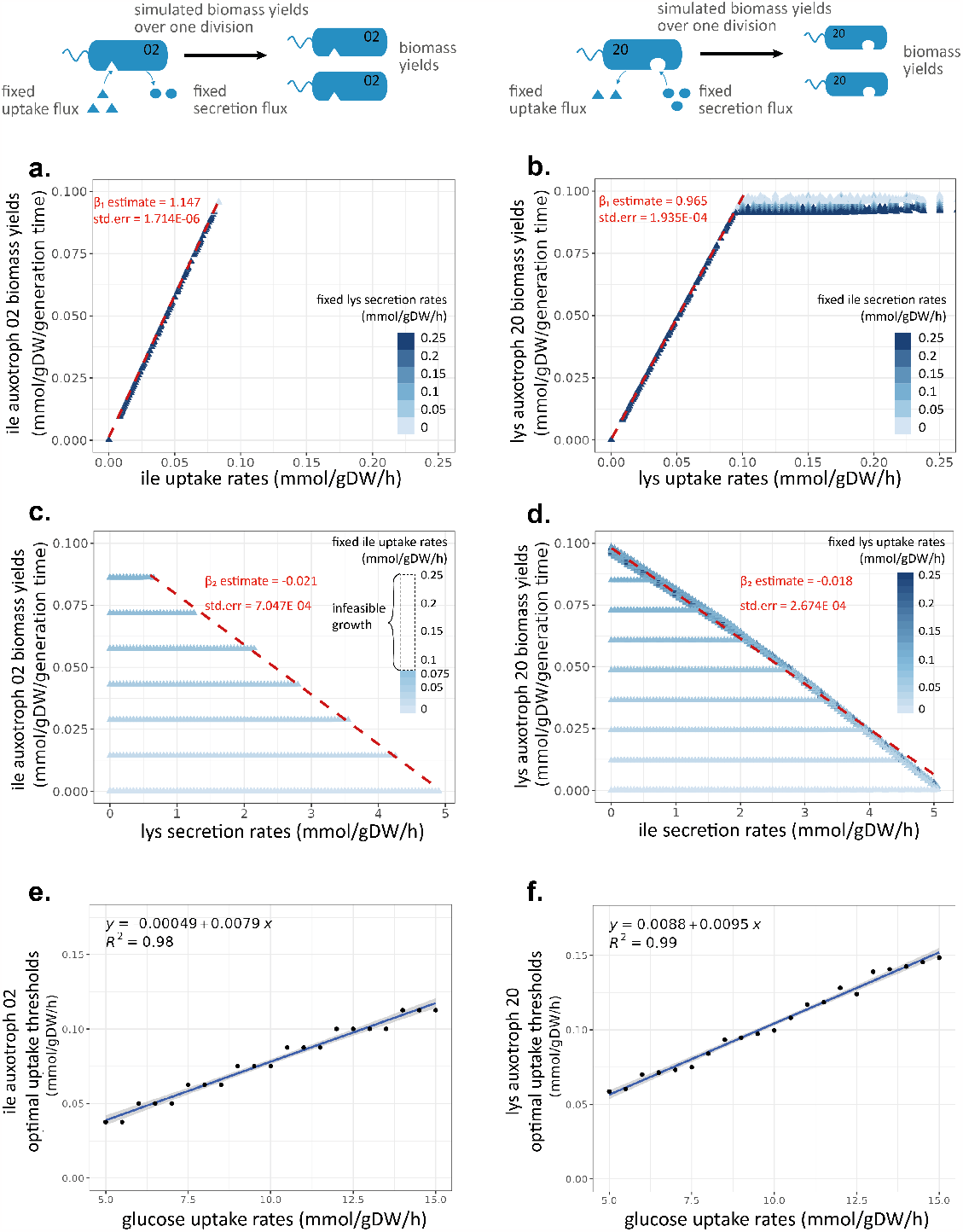
One-generational *ile* / *lys* auxotroph growth simulations when fixing amino acid secretion/uptake rates and increasing uptake/secretion rates. Glucose is used as the sole carbon source ranging from 5 - 15 mmol/gDW/h. **a-b**. One-generational biomass yields of *ile* / *lys* auxotroph strains when fixing *lys* / *ile* secretions and increasing *ile* / *lys* uptake rates from 0 to 0.25 mmol/gDW/h at 0.0125 mmol/gDW/h interval. **c-d**. One-generational biomass yields of *ile* / *lys* auxotroph strains when fixing *ile* / *lys* uptake rates from 0 to 0.25 mmol/gDW/h and increasing *lys* / *ile* secretion rates from 0 to 5 mmol/gDW/h at 0.0125 mmol/gDW/h interval. **e-f**. Linear regressions showing the expected optimal uptake thresholds (OUT) for each unit increase of glucose uptakes by *ile* / *lys* auxotroph strains.

Fig. 2 a-b show one-generational growth results of two auxotroph genotypes by fixing amino acid secretion rates and increasing uptake rates from 0 to 0.25 mmol/gDW/h. Among simulations across glucose supply from 5 to 15 mmol/gDW/h, the biomass yields and amino acid uptake rates of both auxotrophs exhibit a consistent linear relationship, irrespective of glucose uptake levels (Fig. 2 a-b). The linear relationship is disrupted when amino acid uptakes reach certain thresholds (which will be referred to as the Optimal Uptake Thresholds, OUTs). Biomass yields after the OUTs either become unaffected or become infeasible depending on the genotypes.

Fig.2 c-d display similar piecewise growth patterns when fixing amino acid uptake rates at the same range and increasing secretion rates from 0 to 5 mmol/gDW/h. When amino acid uptake rates are below the OUTs, synthesises and secretions of the other amino acid are costless to growth within certain thresholds (which will be referred to as the Costless Secretion Thresholds, CSTs). The negative correlation between CST and amino acid uptake rates suggests interlocking resource allocation processes between amino acid synthesises and uptakes (Fig. 2 c-d).

Different amino acids also display different growth efficiency and secretion costs. Within feasible uptake range and glucose supply at 10 mmol/gDW/h, per unit (0.0125 mmol/gDW/h) increase of amino acid *ile* supports 1.147 unit increase in biomass yields, which is more efficient than the 0.971 increase supported by amino acid *lys* (Fig. 2 a-b). At the same uptake rates of complementary amino acids, *ile* can be secreted over a wider range than *lys* (Fig. 2 c-d).

Based on these observations, the relationships between one-generational biomass yields and amino acid uptake and secretion rates can be portrayed with the following formula:

**Table 1:** Piecewise relationships between one-generational biomass yields and amino acid uptake/secretion rates of *ile* auxotroph 02.

| *ile* auxotroph 02 | $v_{sec} \leq v_{CST}$ | $v_{sec} > v_{CST}$ |
| --- | --- | --- |
| $v_{up} \leq v_{OUT}$ | $Biomass = \beta_1 \cdot v_{up}$ | $v'_{OUT} = v_{OUT} + \beta_2 \cdot (v_{sec} - v_{CST})$<br>$Biomass = \beta_1 \cdot \min\{v_{up}, v'_{OUT}\}$ |

**Table 2:** Piecewise relationships between one-generational biomass yields and amino acid uptake/secretion rates of *lys* auxotroph 20.

| *lys* auxotroph 20 | $v_{sec} \leq v_{CST}$ | $v_{sec} > v_{CST}$ |
| --- | --- | --- |
| $v_{up} \leq v_{OUT}$ | $Biomass = \beta_1 \cdot v_{up}$ | $Biomass = \beta_1 \cdot v_{up} + \beta_2 \cdot v_{sec}$ |
| $v_{up} > v_{OUT}$ | $Biomass = \beta_1 \cdot v_{OUT} + \beta_2 \cdot v_{sec}$ | |

Where *v*_*up*_ and *v*_*sec*_ are the variating amino acid uptake and secretion rates, *β*_1_ represents the linear relationship between biomass growth and amino acid uptakes, *β*_2_ represents the linear growth cost of amino acid secretion after CST, *v*_*OUT*_ is the OUT that vary by genotypes, 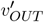 is the reduced OUT of genotype 02 due to high amino acid secretion cost, and *v*_*CST*_ is the value of CST. Parameter *v*_*OUT*_ and *v*_*CST*_ can be obtained from each sample using the following formula:

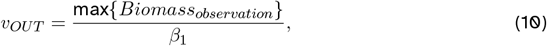

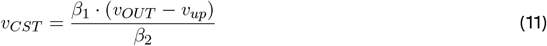

Among the model predictors, both *β*_1_ and *β*_2_ are consistent across glucose uptake levels (Fig. 2 a-b; also see model results in Supplementary Table T1). By contrast, correlation tests between *v*_*OUT*_ and observed glucose uptake rates indicate significant correlations (Fig. 2 e-f). It implies that for each genotype, the same amino acid degradation pathways are used across glucose uptake levels, and their growth benefits are utilised proportionately with glucose uptakes. At the same glucose availability, the studied auxotrophs have a fixed amino acid uptake demand regardless of extracellular amino acid availability.

### Cheating mutants promote higher metabolite exchange intensities over the Optimal Uptake Thresholds

Based on an understanding of how one-generational fitness of bacterial auxotrophs is mediated by metabolite exchange rates, I then simulated multi-generational co-culture growth under various forced amino acid secretion rates and unconstrained amino acid uptake rates with initial glucose supply at 10 mmol/gDW/h (Fig.3 a-c & Supplementary Material S1). By the end of glucose depletion, the cross-feeding resident communities are always the fittest at metabolite exchange intensity 0.1 mmol/gDW/h (Fig. 3 a-b). This intensity is close to but slightly higher than the intensity where auxotrophs have the smallest growth rate differences in monocultures (Fig. 3 d). It is also interesting that their remaining fractions are comparatively high at 1 mmol/gDW/h metabolite exchange intensity (Fig. 3 c), despite the higher growth rate difference in one-generational growth (Fig.3 f). By comparing to auxotrophs’ mono-culture growth with the same secretion and uptake rates of amino acids as the cross-feeding encounters (Fig. 3 e-f), I find a constantly lower growth rate difference in co-culture growth and reversed fitness advantage of cross-feeding auxotroph 02 to 20 during glucose depletion. It indicates that co-existence with cheating mutants reduces fitness difference between cross-feeding strains and increasingly favors strain 02 over 20, which promotes alternate scenarios of cross-feeder survival despite high metabolite exchange costs.

**Figure 3:**
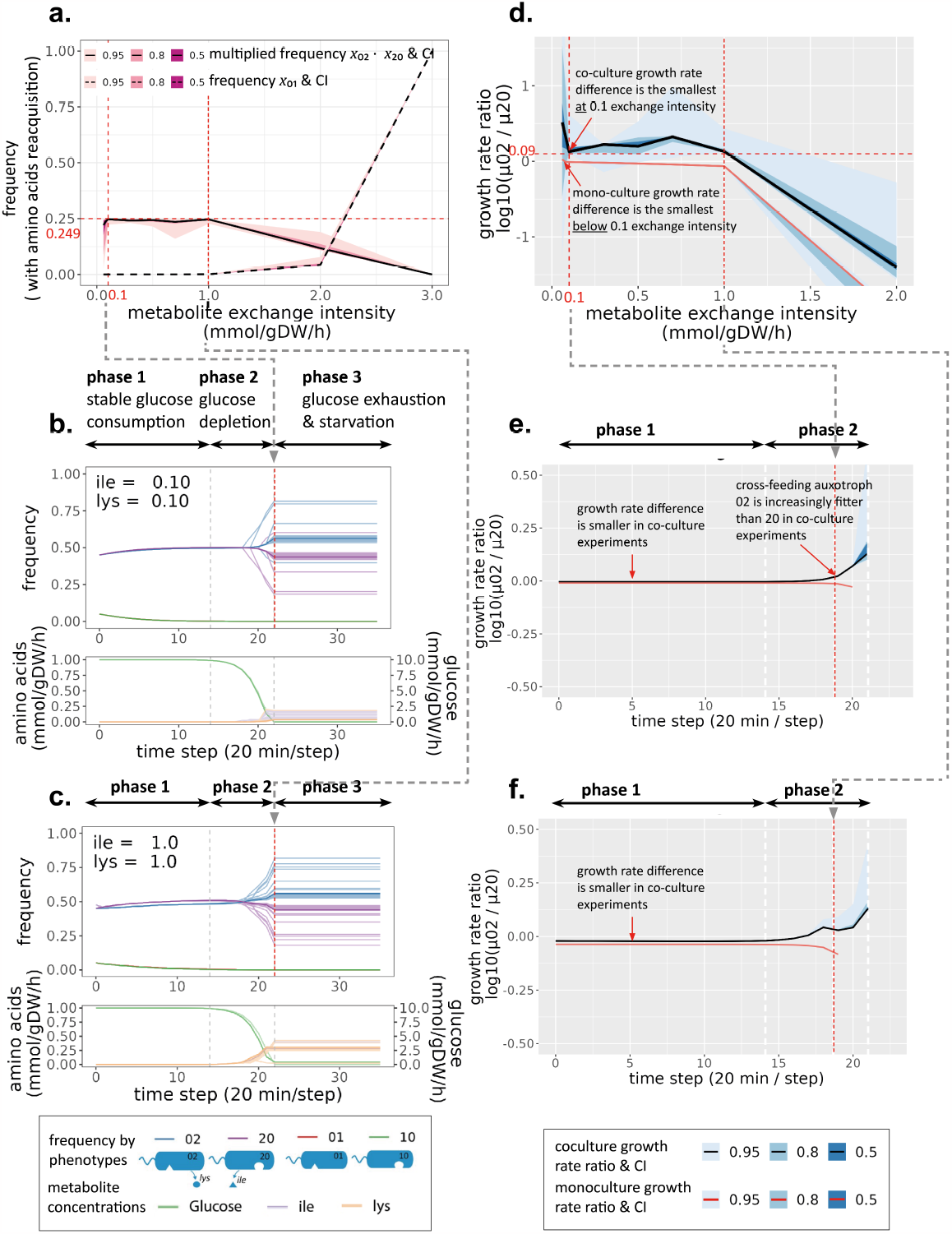
Invasion experiment results including amino acid reacquisition. In each invasion experiment, cross-feeding strains [02,20] are the resident community and cheating mutant strains [01,10] are invaders. Each co-culture grows over 12 hours (generation updated every 20 minutes) with 100 replicates (displayed as faded). **a**. Surviving frequencies of cross-feeders and dominant cheaters (strain 01) by the end of glucose depletion. Multiplied frequency of strain 02 and 20 is used to depict co-existence quality of cross-feeding auxotrophs. **b**. Invasion experiments at metabolite exchange intensity 0.1 mmol/gDW/h, where cross-feeding strains have the highest surviving frequencies. **c**. Invasion experiments at metabolite exchange intensity 1 mmol/gDW/h. Cross-feeding strains have high multiplied frequencies despite high amino acid secretion costs. **d**. Growth rate ratio of cross-feeding auxotrophs 02 to 20 in co-culture and mono-culture experiments before glucose exhaustion. Auxotrophs secrete amino acids at the same rates in both experiments, and receive constant amino acid uptakes without sharing with cheaters in mono-culture experiments. **e-f**. Growth rate ratio of cross-feeding auxotrophs 02 to 20 in co-culture and mono-culture experiments at metabolite exchange intensity 0.1 and 1 mmol/gDW/h.

### Amino acid reacquisition expands the feasible metabolite exchange range at the cost of cross-feeder interdependencies

The invasion experiments discussed above update amino acid uptake rates with additional amino acid surplus. Under this condition, cross-feeding can be maintained as the optimal strategy when metabolite exchange intensity is below 2 mmol/gDW/h (Fig. 3 a). However, crossfeeding can only be maintained under metabolite exchange rates at 0.7 mmol/gDW/h after I blocked amino acid reacquisitions (Fig. 4 a-b). It suggests that amino acid accumulation and reacquisition support cross-feeders’ survival as metabolite secretion costs build up.

**Figure 4:**
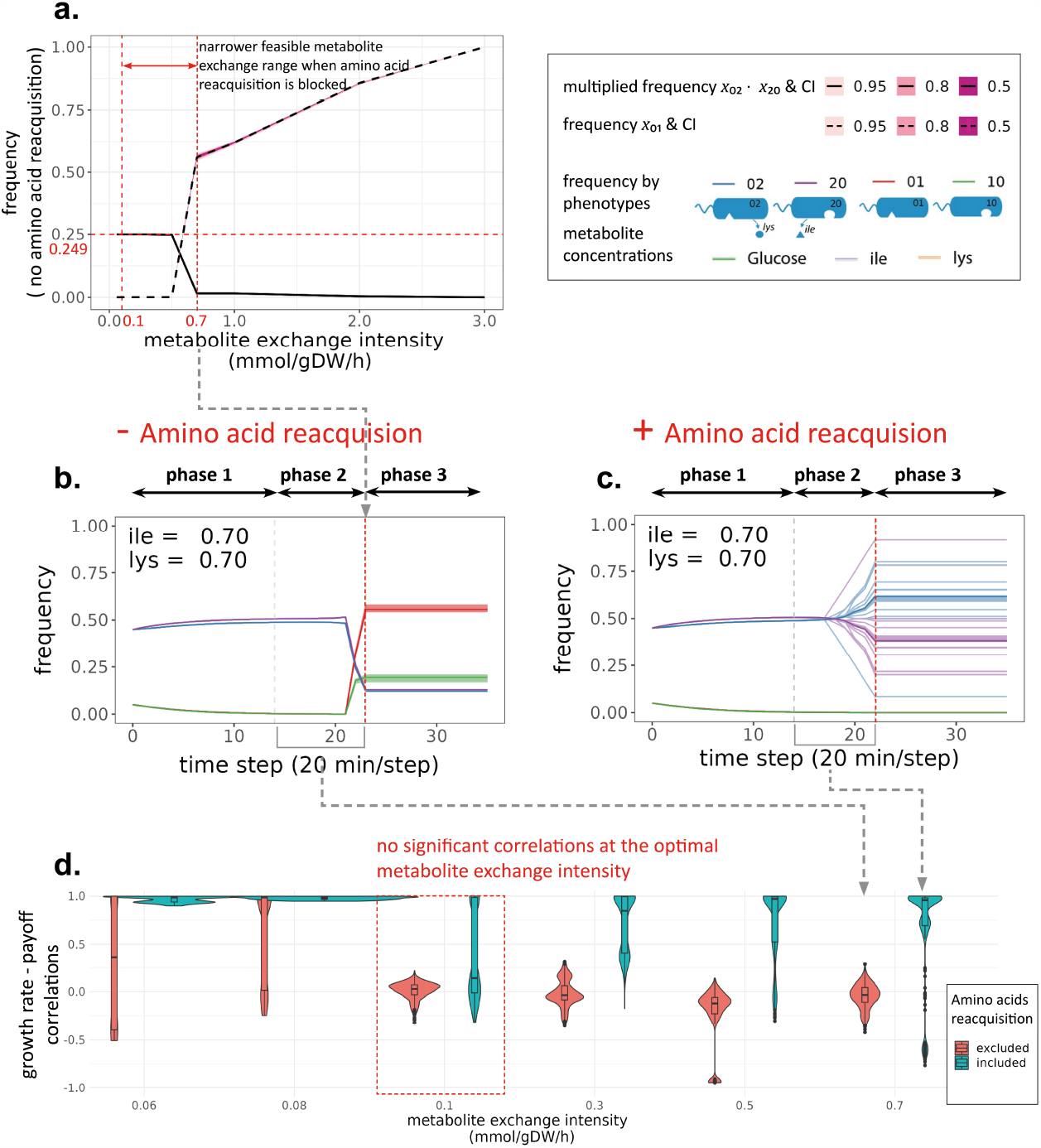
Invasion experiment results excluding amino acid reacquisition. Only glucose concentration is updated to the strain’s uptake rates for each iteration. All other criteria are the same as invasion experiments with amino acid reacquisition. **a**. Surviving frequencies of crossfeeders and dominant cheaters (strain 01) when amino acid reacquisition is blocked. Multiplied frequency of strain 02 and 20 is used to depict co-existence quality of cross-feeding auxotrophs. **b**. Invasion experiments at metabolite exchange intensity 0.7 mmol/gDW/h when amino acid reacquisition is blocked. Cross-feeding auxotrophs are successfully invaded by cheating mutant 01. **c**. Invasion experiments at metabolite exchange intensity 0.7 mmol/gDW/h when amino acid reacquisition is allowed. Cross-feeding auxotrophs co-exist in all replicates by the end of glucose depletion. **d**. Correlation distributions between growth rates of the fittest strain and its payoff advantages over the other strains. Datasets for correlation tests are sampled (30%) from co-culture replicates during glucose depletion phase. Separate correlation tests are conducted for each time step to minimise impacts from payoffs accumulated by time.

On the other hand, amino acid reacquisition significantly enhances coupling between frequency of the fittest strain and its payoff advantages during glucose depletion phase (Fig. 4 d). Except for metabolite exchange near the OUT, the fitter cross-feeding strains at all other metabolite exchange levels are increasingly fitter as it reacquire amino acid surplus. The high coupling explains the cross-feeder survival at 0.7 mmol/gDW/h metabolite exchange intensity (Fig. 4 c) as well as lower multiplied frequencies below 0.7 mmol/gDW/h when amino acid reacquisiont exists (compared between Fig. 4 a and Fig. 3 a). Therefore, amino acid reacquisition not only enhances fitness advantages of cross-feeders over cheaters, but also break the frequencydependent balance between the cross-feeding strains. Fortunately, such disturbance is comparatively small when metabolite exchange intensity is close to and slightly above the OUT (Fig. 4 d).

## 4 Discussion

Predicting and maintaining cross-feeding interactions in a bacterial community is difficult particularly because the metabolite exchange rates and resulting growth rates are sensitive to metabolic environment and difficult to track or control. In this study, I tackled this problem from an evolutionary perspective, integrating evolutionary game theory dynamics and physiological FBA to identify the fittest metabolite exchange rates in a cross-feeding auxotroph community. I discovered that individual fitness of studied auxotrophs is constrained by optimal amino acid uptake thresholds (OUT), which is proportionate to glucose uptake rates. These amino acid uptake thresholds are close to but slightly lower than the fittest metabolite exchange intensity in multi-generational co-culture growth. In addition, reacquisition of accumulated amino acids enhances survival of cross-feeding auxotrophs at high metabolite exchange intensities, but also amplifies fitness differences during glucose depletion phase. Fortunately, the disturbance effect does not significantly hinder synthrophic community success near the OUTs.

In the following sections, I will discuss the revealed mechanisms of fitness trade-offs in metabolite exchange by *E*.*coli* auxotroph strains, how they are regulated by media conditions, and implications for maintaining cross-feeding strategies of engineered syntrophic communities.

### Fitness trade-offs in amino acid exchange

Amino acid isoleucine (*ile*) and lysine (*lys*) are essential building blocks for proteins in bacterial cells as well as key carbon sources^42^. At the same time, glucose provides more accessible carbon atoms and key precursors for various cell components. This is the first study that investigates relationships between different combinations of amino acid uptake / secretion rates and amino acid auxotroph growth rates under various glucose supply levels. The amino acid uptake thresholds (OUTs) I identified have empirical supports from other studies^42–44^, both suggesting that bacterial cells tend to keep amino acid uptakes in a narrow range in order to maintain their optimal homeostasis states for protein productions. Therefore, amino acid uptakes at the identified OUTs will bring the highest growth benefits for a corresponding auxotroph.

In addition to the OUTs, this study finds new relationships between single amino acid uptakes of auxotrophs and the glucose supply. In FBA simulations, amino acid *ile* / *lys* and glucose proportionately contribute to biomass yields of *ile* / *lys* auxotrophs (Fig. 2 a-b). Excessive amino acid uptakes cannot make up for the lack of glucose supply. It suggests that the OUTs of *ile* / *lys* auxotrophs are essentially mediated by glucose supply levels. Reversely, the ideal glucose supply for sustaining *ile* / *lys* auxotrophs can be determined by the desired OUTs.

Before both auxotrophs reaching OUTs, there exists a novel threshold (CST) where amino acid secretions are costless to growth. It means that both auxotrophs have non-overlapping resource allocations between cell growth and amino acid secretions under OUTs and do not need to tradeoff their growth for amino acid cross-feeding below CSTs (Fig. 2 c-d). The existence of CSTs make auxotroph cross-feeding towards their OUTs a win-win strategy to perform.

In summary, how much fitness trade-offs an auxotroph tend to commit is determined by the OUTs of cross-feeding pairs, and mediated by amino acid secretion costs after they reach their own OUT. This process is essentially controlled by initial glucose supply levels, which means the optimal metabolite exchange strategy is highly related to initial glucose availability.

### The optimal metabolite exchange strategy under the presence of cheating mutants

The piecewise costs and benefits of amino acid uptakes and secretions determine auxotrophs’ intrinsic growth rates during one-generational mono-culture growth. In co-culture context, the fitness of these auxotrophs is influenced by other factors such as increasing growth rates of cheaters and accumulated shared metabolites.

With various metabolite exchange intensities, I find the intensity at 0.1 mmol/gDW/h begets the fittest cross-feeding communities, which survive from invasions in all replicates with low variations of final fitness differences (Fig.3 b). Its proximity to OUTs can be explained by fitness trade-offs for win-win status as discussed above. However, this optimal metabolite intensity is not the same intensity that induces the smallest growth rate differences in mono-cultures (Fig. 3 d). As amino acid accumulation effects do not change this fittest intensity value, this is likely the consequence of encountering cheating mutants. In other words, an increase in cheating mutant encounters tend to stimulate higher metabolite exchange intensities among cross-feeding auxotrophs.

Co-culture simulations also show an alternative fitness trade-off scenario for cross-feeder coexistence. Under high metabolite exchange intensities (e.g. 1 mmol/gDW/h in Fig. 3 c), existence of cheaters biasedly selects for *ile* auxotroph 02. As *ile* auxotroph 02 is initially less fit than *lys* auxotroph 20, the reversed fitness advantage during cheaters’ growth makes crossfeeder / cheater co-existence possible with low variations. Such advantage reversal can be used as a strategy for designing resilient cross-feeding auxotroph communities.

By excluding amino acid reacquisition effects in co-culture simulations, I find that cross-feeding auxotrophs survive under a narrower metabolite exchange intensity range with higher co-existence frequencies (Fig. 4 a). This result suggests the amino acid reacquisition is crucial for maintaining cross-feeding interactions at high metabolite exchange intensities, under the threats of cheating mutants. On the contrary, observations without co-existence with cheating mutants suggest a negative impact of reacquiring cross-feeding nutrients (*NH*^+^, McCully et al. ^45^). Therefore, the key benefits of amino acid reacquisition might lie in its rescue effects for amino acid donors when encountering cheaters.

Nonetheless, amino acids reacquision brings risks for cross-feeding communities via enhancing couplings between strain growth rates and its total fitness payoffs against other strains. This finding provides mechanistic explanations for observations by ^29^ when additional amino acids are supplied, cross-feeding auxotrophs with impaired growth rates are less fit than their prototroph ancestors since the growth rates are more coupled with fitness payoffs. Notably, such growth rate payoff couplings are not significant at the identified fittest metabolite exchange intensity (Fig. 4 d). It implies strong frequency-dependent balance still exist between the cross-feeding strains near this metabolite exchange intensity, and reaffirm the high fitness of cross-feeding communities when they exchange amino acids close to and higher than their OUTs.

Bringing together these findings, the fittest metabolite exchange strategy of a cross-feeding community is determined by both initial glucose supply levels and proportion of cheating mutants. A stable cross-feeding community under higher glucose availability or higher proportion of cheating mutants will have higher metabolite exchange intensities. In addition, an environment supportive for amino acid reacquisitions expands the feasible metabolite exchange range when cheaters are present. However, the environment without reacquisition opportunities (e.g. high diffusion rates) can better maintain cross-feeding interactions, as long as the exchange intensities are within the feasible narrowed range.

With my model framework, the evolutionary optimal metabolite exchange intensities can be estimated based on known glucose supply and cheating mutation rates. The value obtained can be used as reference for the metabolite leakage / exchange parameters in multi-species population / metabolism models^4–6^. Reversely, the desired metabolite exchange intensities can instruct experimental design of glucose supply and diffusion speeds for the best maintenance of cross-feeding communities.

### Limitations and future work

Despite these intriguing findings, there are clearly certain caveats in the framework employed in this study. For instance, adaptation of strategies is not considered during invasion processes. As bacteria are adaptive to invasions^46^ as well as changes in resource, they may change the metabolite secretion rates as cheaters gaining advantages and glucose availability drops. Current invasion experiments represent such strategy choice by simulating one strategy with one mutant strategy in each invasion event (Fig. 1), and future work can expand this strategy space in each event, allowing strategy shifts during invasions. Another key simplification underlying my work is the assumption that strains exchange metabolites without much loss in random cellcell encounters. Although efficient physical attachments are common ways of cross-feeding among bacteria^22^, they are not always necessary. Further considerations are required of more realistic encounters and structuring in space^47^, and loss of metabolites in the process. In addition, the metabolite exchange intensities for invasion experiments are coarse in resolution. More finely grained metabolite exchange combinations need to be conducted in order to obtain more accurate optimal intensities. Lastly, I only involve two strains with pairwise interactions in this study, while interactions with three or more strains involve higher order interactions^48^ and fundamentally change the rules for evolutionary game theory dynamics.

## Acknowledgements

We thank Vitor Ferreira, Aditi Madkaikar and Shuixin Li for valuable discussions. We are grateful to the Imperial College London’s Research Computing Service (RCS) team for providing HPC services and supports. The genome-scale metabolic model of E.coli strain is downloaded from the public database of BiGG Models (http://bigg.ucsd.edu/).

