## Supplementary Information for "Deciphering fitness trade-offs in metabolite exchange at the origin of a bacterial cross-feeding community"

**Table T1: Linear mixed model results for *ile / lys* auxotroph growth using the following grouping criteria and formula:**

**groups**

(A) By each amino acid secretion rate in each glucose supply level

(B) By each glucose supply level

**formula**

(1) lmer(growth ~ uptake + (uptake | groupA), growth_data)

(2) lmer(growth ~ 1 + secretion + (secretion | groupB), growth_data)

| Dataset | Predictor | Estimate | StdErr | t | p | formula |
|---|---|---|---|---|---|---|
| *ile* auxotroph growth | *ile* uptake rates | 1.147 | 1.714E-06 | 669242.3 | < 0.001 | (1) |
| *lys* auxotroph growth | *lys* uptake rates | 0.965 | 1.935E-04 | 4990.3 | < 0.001 | (1) |
| *ile* auxotroph growth | *lys* secretion rates | -0.021 | 7.047E-04 | -29.87 | < 0.001 | (2) |
| *lys* auxotroph growth | *ile* secretion rates | -0.018 | 2.674E-04 | -67.93 | < 0.001 | (2) |

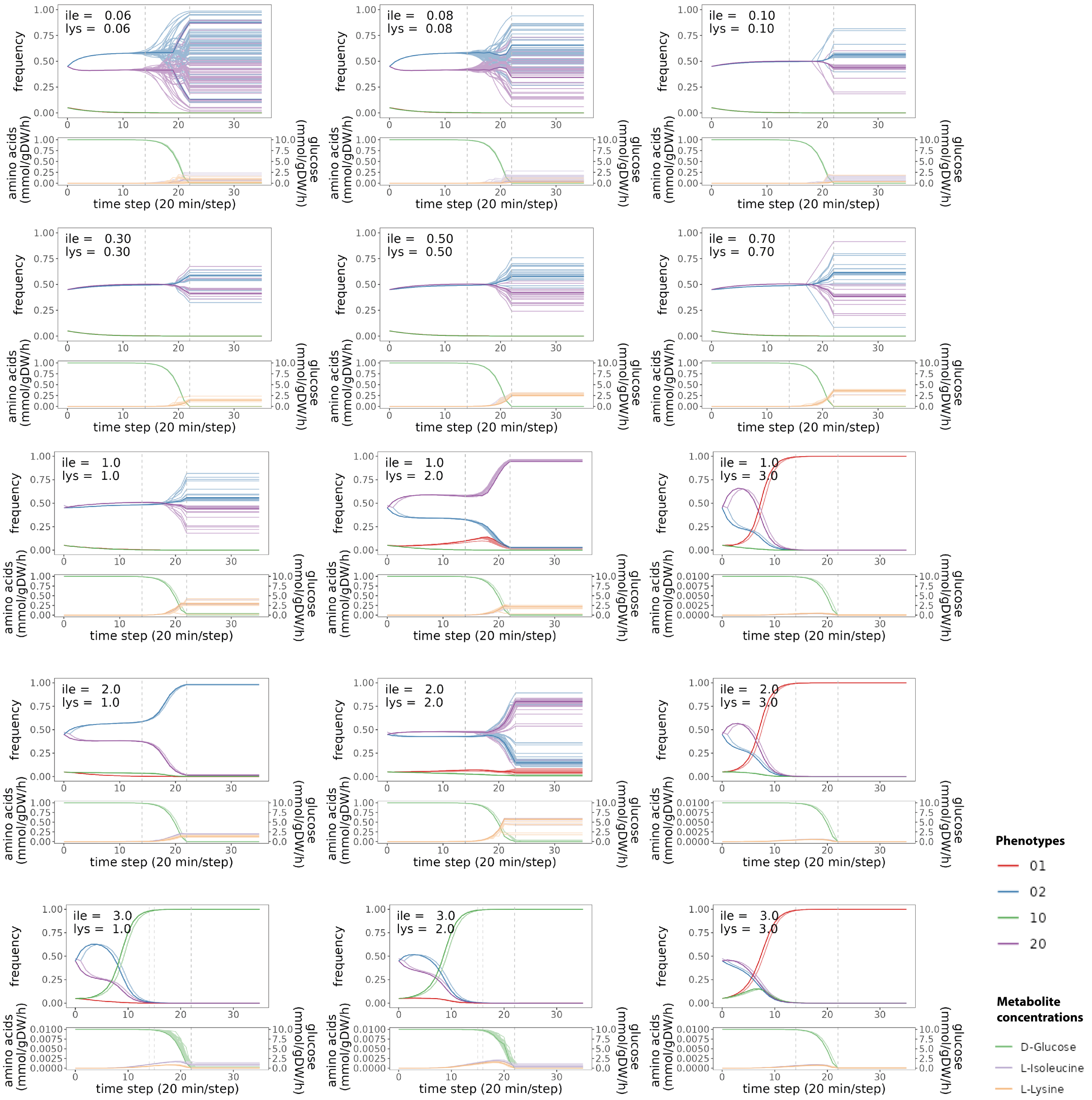


**Figure S1: Supplementary information on all invasion experiments conducted at different amino acid secretion levels.** All invasion experiments with cross-feeding strains [02,20] as the resident community and mutant strains [01,10] as invaders over 12 hours (generation updated every 20 minutes), 100 replicates are conducted and displayed as faded. Metabolite concentrations of glucose, L-Isoleucine (*ile* ), and L-Lysine (*lys* ) are updated and added to constrain maximum uptake rates for each iteration.

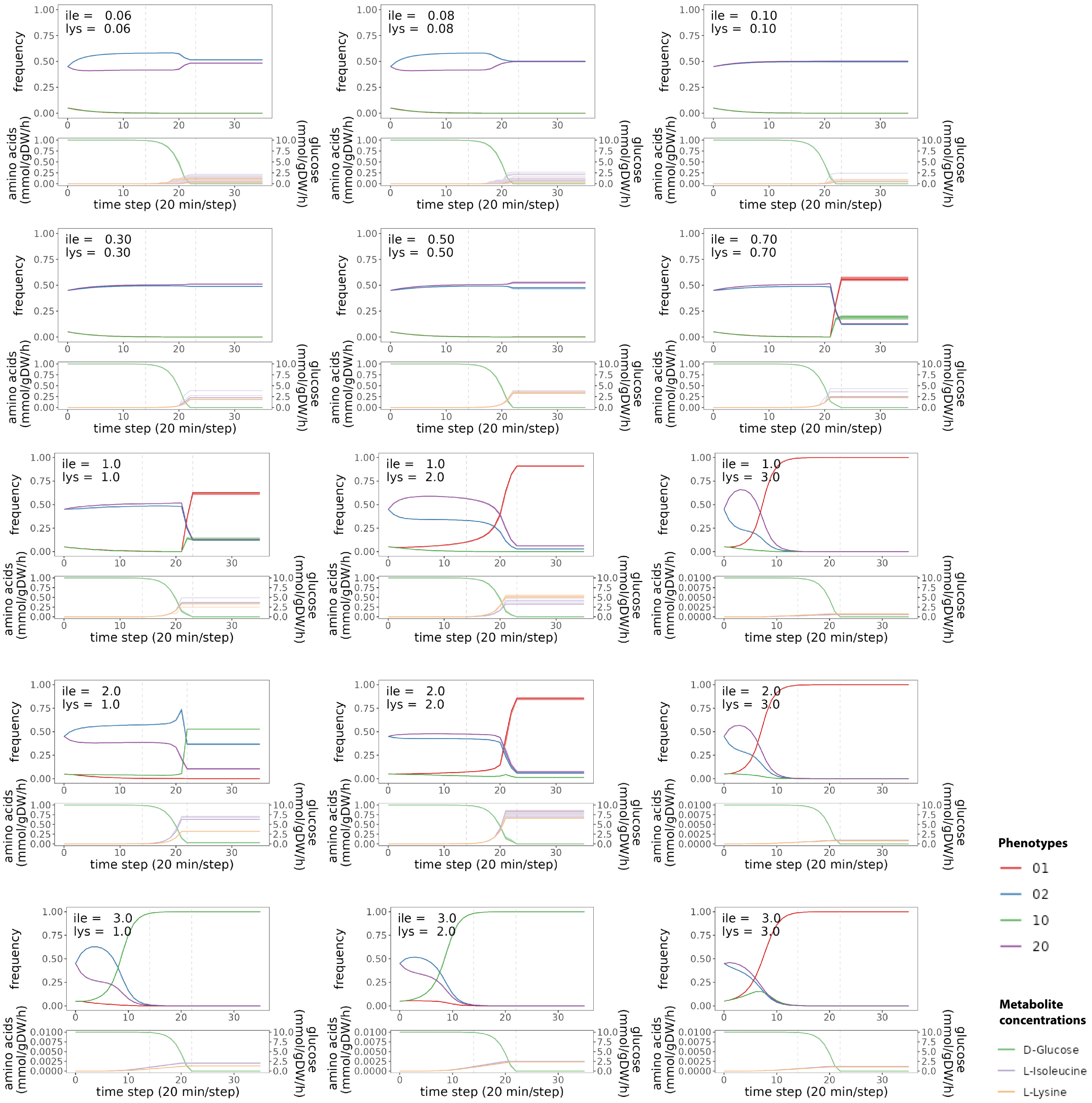


**Figure S2: Supplementary information on all invasion experiments excluding amino acid reacquisition.** Only glucose concentration is added to the strain's maximum uptake rates for each iteration. Other criteria are the same as invasion experiments with uptakes of amino acid surplus, 100 replicates are conducted and displayed as faded.
